# Behavioral Economic Profiles in Theft Recidivists with and without Kleptomania

**DOI:** 10.64898/2026.07.26.740843

**Authors:** Yukiori Goto, Satoshi Yoshino, Chikara Kita, Moojun Won, Young-A Lee

**Affiliations:** Graduate School of Informatics, Kyoto University, Kyoto, 606-8501, Japan; Training Institute for Correctional Officials, Ministry of Justice, Tokyo, 196-0035, Japan; Shogakukan Academy Co., Ltd., Tokyo, 101-0051, Japan; MRC Lab Clinics, Tokyo, 181-0012, Japan; Department of Food Science and Nutrition, Daegu Catholic University, Gyeongsan, 38430, Korea

**Keywords:** Neurocriminology, Kleptomania, Addiction, Risk/Loss aversion, Prefrontal cortex

## Abstract

Recurrent theft, such as shoplifting, is a devastating social problem that poses a significant economic burden on society. However, the neurobehavioral mechanisms underlying such criminal offenses remain barely understood. Here we investigated the behavioral economic profiles of theft recidivists (TR) with and without kleptomania. To identify the decision-making processes underlying stealing behaviors, the hypothetical purchase task and the balloon analogue risk task (BART) structured with gain and loss frames were administered, along with concurrent measurement of prefrontal cortex (PFC) activity during the BART using functional near-infrared spectroscopy. TR with kleptomania demonstrated heightened loss aversion and price sensitivity (elasticity) compared to TR without kleptomania and control subjects with no criminal records, with a significant correlation between these factors, suggesting that heightened loss aversion may contribute to higher elasticity. Moreover, the relative loss-to-risk aversion ratio was found to be positively and negatively associated with the right and left PFC, respectively, suggesting that an imbalance between left and right PFC activity may be involved in the altered behavioral economic profiles. These findings challenge the conventional view of theft as a reward-seeking behavior, revealing that an abnormally inflated fear of monetary loss may play a role in recurrent stealing in kleptomania.

## INTRODUCTION

Theft recidivism is a major social issue that imposes a substantial economic burden on society. For instance, the financial loss caused by shoplifting in the US has been estimated to be 47.8 billion dollars in 2025 (Shopping, 2026). The rate of theft recidivism is also estimated to be even higher than 70% within a 5-year period if no intervention takes place (National Academies of Sciences, 2020). Despite such a devastating social impact, the neurobehavioral mechanisms underlying property offenses remain largely unknown. A significant number of cases of recurrent thefts could also be associated with psychiatric conditions, most notably kleptomania, an impulse control disorder, yet recently suggested to share features with addiction disorders (Grant, 2006). Kleptomania was initially identified over 200 years ago; since then, however, few comprehensive studies have explored its exact neurobehavioral mechanisms to date (Asaoka, Won, Lee, & Goto, 2025).

Behavioral economics offers a framework for understanding decision-making processes, particularly how individuals evaluate potential gains and losses, based on the principles of risk and loss aversion and framing effects (Tversky & Kahneman, 1981). These economic decision-making processes have been evaluated using various behavioral tasks, including the hypothetical purchase task (HPT), which is widely used to measure operant demand and price sensitivity (Roma, Reed, DiGennaro Reed, & Hursh, 2017), and the balloon analogue risk task (BART), particularly when structured with framing effects for gain and loss frames, serves as a metric for quantifying risk and loss aversion (Schulman, Chong, & Lockenhoff, 2022; Xu, Wang, Liu, Wang, & Zhang, 2020). Studies have shown that the neural mechanisms underlying risk and loss aversion are associated with the prefrontal cortex (PFC). Thus, accumulating evidence suggests that the left and right hemispheres of the PFC play asymmetric roles in mediating risk and loss aversion, with the left PFC tied to risk aversion for evaluating positive outcomes and promoting deliberate caution, whereas the right PFC is engaged in loss aversion for processing negative outcomes and threat assessment (Huang et al., 2017; Ye, Chen, Huang, Wang, & Luo, 2015).

In the context of psychiatric disorders, behavioral economic profiling has been extensively used to study addictive disorders. For instance, addiction studies utilizing the HPT have reported that the severity of addiction symptoms in affected individuals can be predicted from indices representing high and inelastic demand in the task (Berry, Naude, Johnson, & Johnson, 2023; Gonzalez-Roz, Martinez-Loredo, Secades-Villa, Amlung, & MacKillop, 2020; Strickland & Lacy, 2020). The literature also indicates that individuals with substance abuse exhibit increased risk-taking, which predicts symptom severity (Canning, Schallert, & Larimer, 2022), whereas other studies have demonstrated decreased loss aversion in these individuals (Genauck et al., 2017; Thrailkill, DeSarno, & Higgins, 2022), collectively suggesting that addition disorders may involve an imbalance of higher risk-taking and decreased loss aversion (Krmpotich et al., 2015). However, it remains unknown whether theft recidivists, including those with kleptomania, share addiction-like profiles.

Given the lack of insight into economic decision-making processes associated with theft recidivism, this study aimed to investigate alterations in the behavioral economic characteristics of theft recidivists, specifically differentiating between those with and without kleptomania. Based on the phenotypic similarities between recurrent stealing and addictions, we hypothesized that theft recidivists would exhibit greater risk-taking and reduced loss aversion, along with altered PFC activity, mirroring the decision-making profiles of individuals with addiction disorders. This hypothesis was tested using the HPT and BART with framing effects to evaluate the risk and loss aversion. While the participants performed the BART, PFC activity was sampled using functional near-infrared spectroscopy (fNIRS).

## METHODS

### Subjects and Methods

A total of 63 theft recidivists (TR) and 53 control subjects (CT) were recruited for this investigation, along with other studies. The sample size was determined to achieve 80% power with a moderate to large effect size for group comparison. The inclusion criteria for the TR group were prior records of incarceration due to larceny, primarily shoplifting, between 18 and 79 years old, and living in Japan at the time of the investigation, whereas the inclusion criteria for the CT group were no criminal records at the age of 18-79 years old living in Japan. The exclusion criteria were the inability to understand the investigation details owing to intellectual disability or any other reasons, and none met the exclusion criteria. However, seven TR participants dropped out from the investigation due to loss of interest in participation, and one CT participant was removed from the data analysis since they engaged in the task without understanding the procedure of the BART. Sixteen TR participants were diagnosed with kleptomania (TR-KA) and were receiving treatment at clinics at the time of the investigation. Socioeconomic status was not considered for inclusion/exclusion criteria, as all TR recruited for the investigation had been in rehabilitation centers or welfare facilities for their reintegration into society, such that their socioeconomic status is relatively invariable.

### Hypothetical Purchase Task (HPT)

The HPT (Roma et al., 2017) was administered to examine operant demand, a behavioral economic concept that measures the relationship between the consumption of a reinforcer and the cost of obtaining it (Reed, Hursh, Berry, & Strickland, 2025).

The HPT consists of a series of questions. Participants were first asked to recall any specific snack or sweet that they frequently consumed or liked the most and to provide the unit price. The unit price of the snack/sweet was not different between groups (193.32 ± 9.71 Japanese yen in CT; 195.54 ± 16.16 in TR; 245.63 ± 52.42 in TR-KA; 177.73 ± 11.05 in TR without kleptomania [TR-NK]). Then, the participant was asked to imagine how many of them they would like to purchase in one month (but they must eat them on the same day of obtaining) when the product was sold at a unit price of 0 (free), 1, 5, 20, 50, 200, 600, 1000, and 5000 Japanese yen (JPY; JPY 200 is approximately US$1.5).

The measurements of the HPT included the intensity of demand or unconstrained demand Q_0_ (a quantity of the product purchased at free), the elasticity of demand or price sensitivity α (a rate of change in purchase as the unit price increases; Suppl. Fig. S1), the maximum expenditure O_max_ (a maximal financial allocation for purchase at any unit price), the price at maximum expenditure P_max_ (a unit price at which Omax is achieved), and break point (BP; the first price at which consumption drops to 0).

### Ballon Analogue Risk Task (BART)

The BART with a framing effect (Schulman et al., 2022; Xu et al., 2020) was used to examine risk and loss aversion in this study. The task was programmed and administered using the Inquisit Web (Millisecond Software, LLC).

BART was originally developed to assess risk-taking by counting how much a participant pumps up a balloon presented on a screen to earn money. Thus, in this task (hereby denoted as the gain frame of the BART), a participant pumped up a balloon freely upon presentation, with which the balloon grew, and a small amount of money was added at each pump (JPY 20) to the potential earnings. However, the risk that the balloon burned increased as the participant pumped more, and if the balloon burned, the potential earnings were lost, so that the participant had to stop pumping and collect the earnings before starting the next trial (another balloon pumping). Participants were asked to earn as much money as possible. The task consisted of 20 trials (20 balloons).

In addition to the grain frame of the BART, another version of the BART (denoted as the loss frame of the BART) was administered to participants. In this version, JPY6,000 (approximately US$40) was given to a participant first, and JPY300 (approximately US$2) was subtracted upon presentation of a balloon. Thus, 20 trials (20 balloons) of presentations resulted in zero earnings. However, pumping a balloon decreased the subtraction for a small amount at each pump (JPY 20), so that the participant could save the money in their possession. However, similar to the grain frame, the risk that the balloon burned increased as the participant pumped more, and if the balloon burned, the potential savings were lost.

The gain and loss frames of the BART were administered sequentially to each participant, with counterbalancing the order of presentation. The adjusted average pumps (the number of pumps that a participant made in a trial in which a balloon was not burned) were measured as risk-taking in the gain frame and loss aversion in the loss frame. The log ratio of the adjusted average pumps in the loss frame to those in the gain frame (log(L/R)) was measured for each participant. Thus, it represents a relative loss-to-risk aversion tendency, with a higher ratio indicating more loss aversion and less risk-taking, whereas a lower ratio indicates more risk-taking and less loss aversion.

### fNIRS

PFC responses while the participants performed the BART were measured using fNIRS. Due to logistical and environmental constraints, fNIRS was conducted only in CT and TR-NK, but not TR-KA, participants, while they engaged in the BART with gain and loss frames.

The NIRO-200 NIRS Image Processing and Measuring System (Hamamatsu Photonics K.K.) was used for the measurements. The system consisted of two emitters delivering laser pulses at wavelengths of 775, 810, and 850 nm, and eight detectors with a distance of 3.0 cm between the emitters and detectors. Oxygenated (O2Hb) and deoxygenated (HHb) hemoglobin changes were sampled over time at a sampling rate of 0.5 Hz at 10 locations (denoted as R1‒R10) spanning the left and right hemispheres of the PFC. R4 and R6 covered the left and right rostral dorsal PFC (PFrd), corresponding to Brodmann area (BA) 10/9/8; R5 and R7 covered the left and right caudal dorsomedial PFC (PFcdm), corresponding to BA6/8; R3 and R8 covered the left and right rostral dorsolateral superior PFC (PFrdls), corresponding to BA10/9; R2 and R9 covered the left and right caudal dorsolateral PFC (PFcdl), corresponding to BA45/46/9; and R1 and R10 covered the intermediate zone between PFrdls and PFcdl of the MarsAtlas (Auzias, Coulon, & Brovelli, 2016).

The magnitudes of the O2Hb and HHb signals were quantified as signed root mean squares (sRMS) as follows:

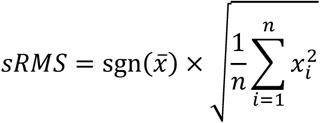

where sgn is a sign function, *x̄* is the mean of the signal, *x*_i_ is the amplitude of the signal at the ith sampling, and n is the total number of samplings. Then, the subtraction of sRMS between the gain and loss frames (L-R sRMS) was calculated at each location such that the L-R sRMS for the O2Hb and HHb signals at each location resulted in 20 independent variables.

### Experimental Design

This study was conducted in accordance with the Declaration of Helsinki and the Ethical Guidelines for Medical and Health Research Involving Human Subjects of the Japanese Ministry of Health, Labour, and Welfare. All procedures were approved by the Human Research Ethics Committee of the Kyoto University Graduate School of Informatics. Written informed consent was obtained from all participants before the investigation. Age, sex, and smoking status were collected at the time of enrollment in the study. The HPT was administered first, followed by the BART with gain and loss frames, along with fNIRS recordings. Due to environmental and logistical constraints, fNIRS could not be conducted in TR subjects who had been diagnosed with kleptomania.

### Data Analysis

All data are expressed as mean ± standard error of the mean (s.e.m.). Statistical significance was set at p < 0.05.

*Analysis of covariance (ANCOVA):* A comparison of dependent measurements between groups was conducted using ANCOVA, along with adjustment for age, sex, and smoking status as covariates. When dependent variables were found to violate normality (Shapiro-Wilk test for normality) and homogeneity (Levene’s test for equality of variance) by assumption checks, transformation of the data using rank-based inverse normal transformation was conducted prior to the statistical analyses. Post-hoc comparisons were conducted using bootstrapped robust t-tests (1,000 bootstraps) with a Bonferroni correction.

*Nonparametric statistical comparison:* Whenever applicable, analyses for comparisons or correlations of two groups were conducted using the nonparametric Mann-Whitney U test and Spearman’s rank order test, respectively.

*Lasso-OLS regression:* Given the overfitting problem of ordinary multiple regression with high-dimensional settings, a two-step Lasso-ordinary least square (OLS) regression approach (Daneshvar & Mousa, 2023; Tibshirani, 1996) was utilized to identify the most robust predictors of log(L/R), the measurement of BART, from the initial set of 20 L-R sRMS variables, the measurements of fNIRS. Prior to the analysis, the predictor variables were standardized. First, feature selection was performed using regularized linear (Lasso) regression. The tuning parameter λ was determined using 10-fold cross-validation. To address the inherent instability of the Lasso variable selection across random data splits, stability selection was conducted by 50 iterations of the tuning parameter selections. A random seed was set prior to each split to ensure the exact reproducibility of the final model. The selection probability at higher than 90%, along with the average of regression coefficient β larger/smaller than ±0.015, of variables were selected for the following unpenalized standard OLS regression with bootstrapping (5,000 straps) to estimate effect sizes. In the OLS, a group factor (CT and TR-NK) was included as a dummy (categorical) variable. Since the same data were used for both selection with Lasso regression and inference with OLS regression, the resulting p-values were exploratory and interpreted as conditional on the selection process rather than as strict tests of universal significance.

*In-silico perturbation analysis:* Since fNIRS samplings were not available for TR-KA, an in-silico perturbation analysis (counterfactual simulation of group differences; (Mana et al., 2023; Sanz Perl et al., 2021)) was conducted utilizing the Lasso regression model derived from CT and TR- NK to investigate whether the log(L/R) difference between TR-KA and the combined CT and TR-NK group could be theoretically explained by variations in the identified fNIRS signals in the PFC. This simulation was predicated on the assumption that the underlying neural architecture mediating BART performance is conserved across both groups. To determine the theoretical PFC activity shift required to bridge the behavioral difference, we calculated the difference between the mean log(L/R) of TR-KA (*Ȳ*_TRKA_) and that of the combined CT and TR-NK group (*Ȳ*_CT/TRNK_). Then, this gap was divided by the sum of the unstandardized β coefficients of the identified PFC areas in the Lasso regression model:

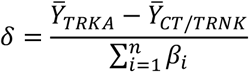

where δ represents the uniform mathematical shift required across all identified signals (n) in the PFC to produce the mean log(L/R) in TR-KA. For biological plausibility assessment of the required shift δ, we evaluated it against the physiological boundaries of sRMS observed in the identified PFC areas of the combined CT and TR-NK group (Min = −448.83, Max = 164.28). Furthermore, to standardize this shift into a comparable metric, δ was divided by the pooled standard deviation (SD = 63.02) of the sRMS of the identified signals in the PFC. This allowed us to express the required shift as a standardized effect size, providing a quantitative measure to determine if the hypothetical variation in PFC activity was within realistic physiological limits.

### Transparency and Openness

We have reported our sample size, data inclusions and exclusions, manipulations, and measures in the study, and that the study followed JARS (Appelbaum et al., 2018). Research materials are available upon request to the corresponding author. Data were analyzed using JASP ver. 0.97.1 (Team, 2026) and OriginPro ver. 2026 (OriginLab Corporation, Northampton, MA, USA), This study’s design and its analysis were not pre-registered.

## RESULTS

Age (Mann-Whitney U test, U = 1483, p = 0.743, Rank-Biserial Correlation r = −0.037) and sex (χ^2^ continuity correction, χ^2^(1) = 0.002, p = 0.968) did not differ between the TR and CT groups, whereas smoking was more prevalent in the TR group than in the CT group (χ^2^(1) = 12.54, p < 0.001).

### Operant Demand

HPT was administered to assess the relationship between the consumption of a reinforcer and the cost of obtaining it in the TR. ANCOVA, adjusted for age, sex, and smoking status, revealed a significant group difference in Q0 (intensity of demand; F(2, 101) = 6.422, p = 0.002, effect size ω^2^ = 0.087; Table 1; Fig. 1a). The post-hoc robust t-test with 1,000 bootstraps and Bonferroni correction found that Q0 was higher in TR with kleptomania (TR-KA) than in CT (t = 2.757, p = 0.021, Cohen’s d = 0.795; Table 1), as well as higher in TR without kleptomania (TR-NK) than in CT (t = 3.105, p = 0.007, d = 0.695; Table 1). However, no difference was observed between the TR-KA and TR-NK groups (t = 0.337, p = 1.000, d = 0.100; Table 1). Moreover, a significant group difference was observed in α (elasticity of demand; F(2, 101) = 4.737, p = 0.011, ω^2^ = 0.060; Table 1; Fig. 1b). In particular, post-hoc tests revealed that α was higher in TR-KA than in both TR-NK (t = −2.940, p = 0.012, d = −0.874) and CT (t = −2.727, p = 0.022, d = −0.786), whereas there was no difference between TR-NK and CT (t = 0.391, p = 1.000, d = 0.088). No group differences were observed in other measurements, such as O_max_ (maximum expenditure; F(2, 101) = 0.645, p = 0.527, effect size ω^2^ = 0.000; Table 1), P_max_ (price at maximum expenditure; F(2, 101) = 2.736, p = 0.069, effect size ω^2^ = 0.029; Table 1), and BP (break point, the first price at which consumption drops to 0; F(2, 101) = 0.931, p = 0.397, effect size ω^2^ = 0.000; Table 1). These results suggest that the baseline craving for the reinforcer at no cost is higher in TR, whereas individuals with kleptomania, but not other TR, exhibit uniquely high price sensitivity for purchase.

**Figure 1.**
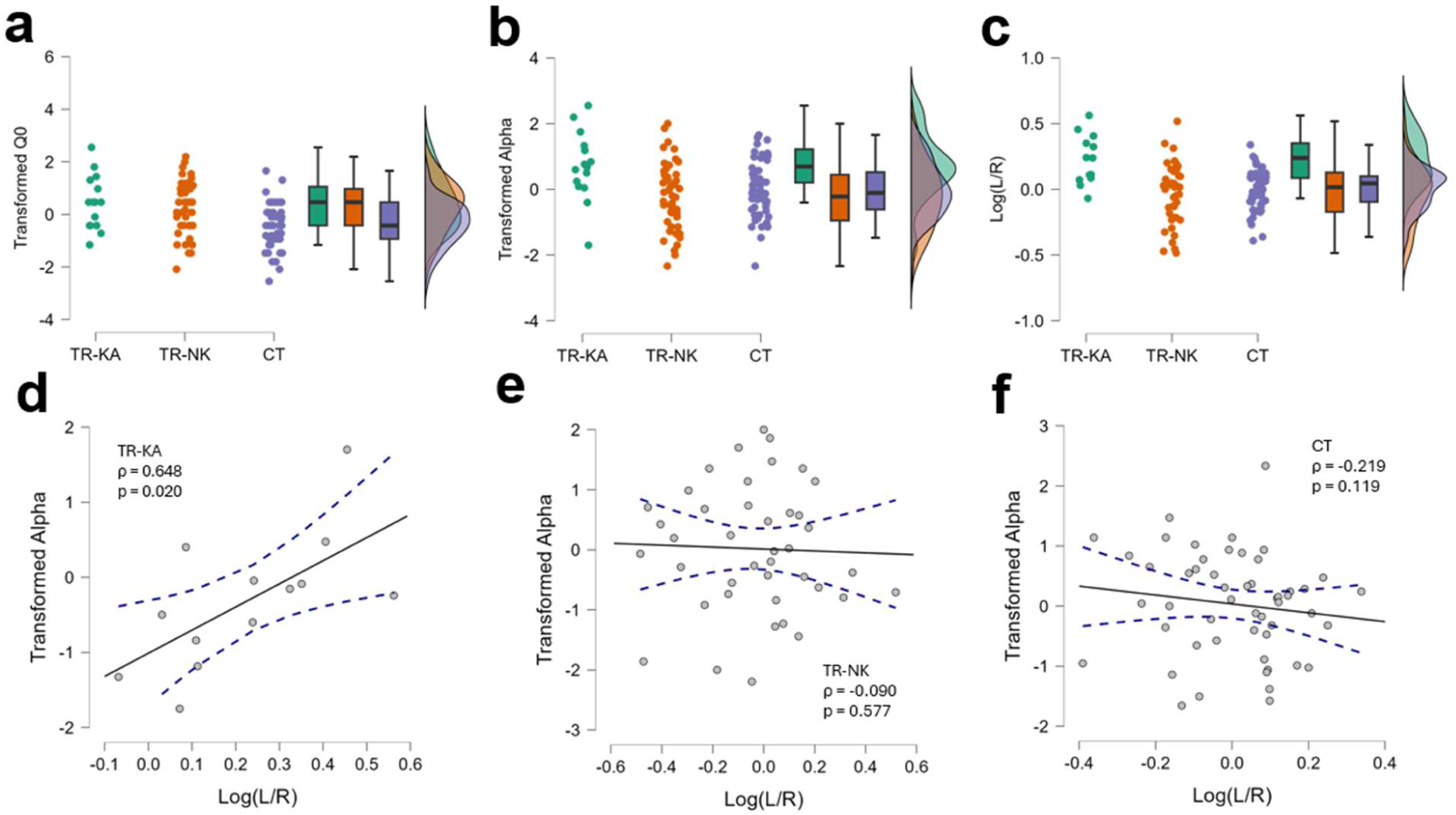
Comparisons of HPT and BART parameters between TR-KA, TR-NK, and CT. a-c, Raincloud plots comparing transformed (data with rank-based inverse normal transformation) Q0 (a) and elasticity α (b) in the HPT and log(T/R), the log ratio of the Loss over the Gain frames in the BART (c) between theft recidivists with kleptomania (TR-KA), those without kleptomania (TR-NK), and participants with no criminal records (CT). **d-f,** Panels showing the correlations between α in HPT and log(T/R) in the BART for TR-KA (d), TR-NK (e), and CT (f). Dashed lines indicate 95% confidence intervals.

**Table 1.** A summary of HPT and BART in CT, KA, and TR.

| Group | HPT |  |  |  | BART |  |
| --- | --- | --- | --- | --- | --- | --- |
| | $Q_0$ | $O_{max}$ | $P_{max}$ | BP | $\alpha$ | Log(L/R) |
| CT | 20.66 ± 5.263 | 1499 ± 202.7 | 348.9 ± 94.80 | 1740 ± 259.4 | 0.446 ± 0.024 | 0.006 ± 0.022 |
| TR-KA | 112.3 ± 62.20 | 5594 ± 3156 | 124.9 ± 38.67 | 1838 ± 476.2 | 0.601 ± 0.053 | 0.225 ± 0.052 |
| TR-NK | 76.35 ± 23.55 | 19660 ± 10310 | 293.8 ± 111.0 | 1694 ± 295.7 | 0.414 ± 0.030 | -0.030 ± 0.035 |

HPT – Hypothetical purchase task; BP – Breakpoint; BART – Ballon analogue risk task; L/R – Ratio of adjusted pump presses in loss over gain frames; CT – Control subjects with no criminal record; TR-KA – Theft recidivists with kleptomania; TR-NK – Theft recidivists without kleptomania. Data represented as mean ± s.e.m.

### Loss/Risk Aversion

To further investigate altered economic decision-making processes in TR, BART was conducted with Gain and Loss frames, with which risk and loss aversions were assessed, respectively, in participants. The log ratio of the adjusted pumps in the loss frame over those in the gain frame (the relative loss-to-risk aversion ratio, log(L/R)) was measured for each subject. Thus, it indicates relative loss/risk aversion tendency, with a higher ratio indicating more loss aversion and less risk taking, whereas a lower ratio indicates more risk taking and less loss aversion. A significant group difference in this ratio was found using ANCOVA (F(2, 101) = 8.870, p < 0.001, effect size ω^2^ = 0.129; Table 1; Fig. 1c). Post-hoc robust t-test revealed that the log(L/R) score was significantly higher in TR-KA than in TR-NK (t = 4.161, p < 0.001, d = 1.356) and CT (t = 3.539, p = 0.002, d = 1.102). No difference was found between the TR-NK and CT groups (t = - 1.113, p = 0.805, d = −0.253). In addition, a striking positive correlation was observed between log(L/R) and α in HPT in TR-KA (ρ = 0.648, p = 0.020; Fig. 1d), but not in TR-NK (ρ = −0.090, p = 0.577; Fig. 1e) and CT (ρ = −0.219, p = 0.119; Fig. 1f).

These results suggest that theft recidivists with kleptomania are characterized by primarily heightened loss aversion (Suppl. Fig. S2), whereas risk/loss aversion in theft recidivists without kleptomania is not different from that in control subjects with no criminal record. In addition, such heightened loss aversion in kleptomania may contribute to higher price sensitivity in the HPT.

### Loss/Risk Aversion-related PFC Responses

To infer the neural mechanisms underlying heightened loss aversion in kleptomania, fNIRS was used to measure PFC responses in 53 CT and 39 TR-NK participants while they engaged in the BART with gain and loss frames, although this could not be conducted directly in TR-KA participants due to constraints.

O2Hb and HHb signals were sampled from 10 locations (R1-R10) over the left and right PFC. The magnitudes of the signals were quantified as signed root mean square (sRMS) and subtracted between the gain and loss frames (L-R sRMS) at each location. Associations between the relative loss-to-risk aversion ratio and L-R sRMS for O2Hb and HHb at each location (a total of 20 variables) were evaluated using the Lasso-OLS regression approach.

Feature selection was performed using Lasso regression, which identified five variables (O2Hb-R7, O2Hb-R8, HHb-R1, HHb-R5, and HHb-R6; λ = 0.016, cross-validated mean square error (CV-MSE) = 0.045, R^2^ = 0.417, at the average with 50 iterations; Fig. 2a, b). These variables were subsequently fitted to an unpenalized standard regression model. With these five variables, a significant model was found (F(6,83) = 3.534, p = 0.004), although its proportion of variance explained was relatively low (R^2^ = 0.203). In particular, negative associations were found in the variables on the left PFC (HHb-R1, unstandardized β = −0.047, p = 0.014; HHb-R5, β = −0.049, p = 0.014), whereas positive associations were observed in the variables on the right PFC (O2Hb-R7, β = 0.041, p = 0.010; O2Hb-R8, β = 0.039, p = 0.014; HHb-R6, β = 0.048, p = 0.008).

**Figure 2.**
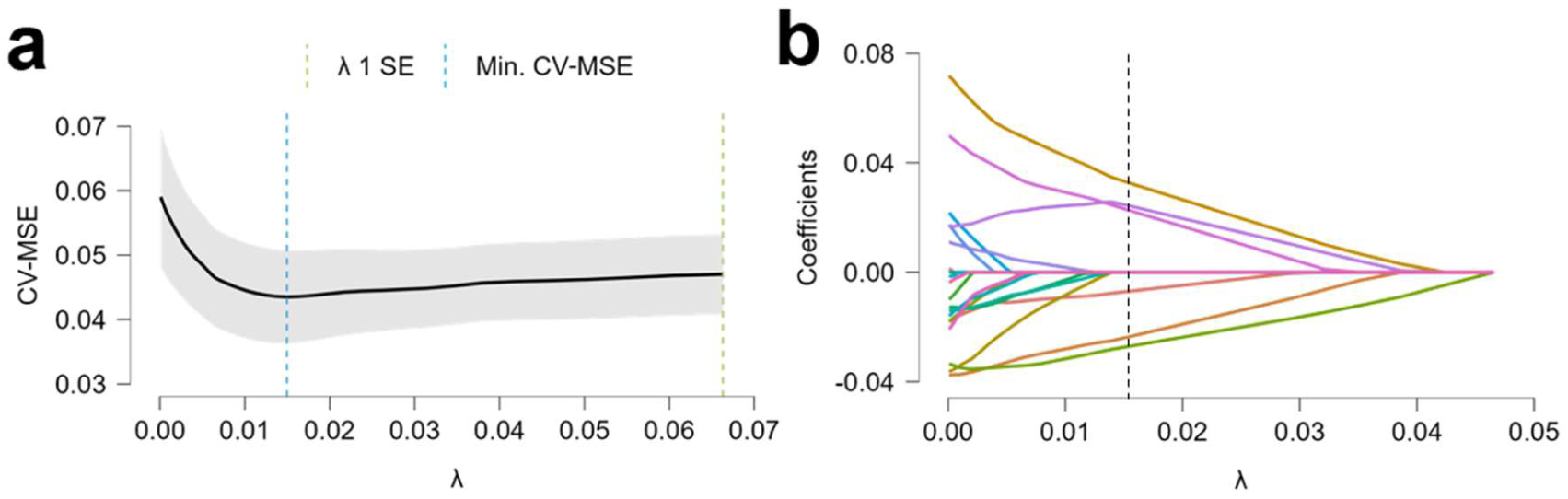
A representative model of Lasso regression. a, Evaluation plot of the penalty parameter λ, showing the overall shape of the error curve, along with the exact point where the error is minimized (blue dashed line) and the point representing the most regularized model within one standard error of the minimum (green dashed line). **b,** Coefficient path (variable trace) plot, illustrating variable importance and selection. The colored lines indicate each variable, and the vertical dashed line indicates the chosen optimal λ.

These results suggest that an imbalance in left and right PFC activity may be associated with altered loss/risk aversion, with stronger left than right PFC activity causing more risk-taking, whereas stronger right than left PFC activity causes more loss aversion.

### PFC Response Simulation for TR-KA

To test the 5 identified signals in the PFC (HHb-R1, HHb-R5, O2Hb-R7, O2Hb-R8, HHb-R6) in the combined CT and TR-NK could explain the higher log(L/R) in TR-KA than CT and TR-NK, we performed a counterfactual simulation using the Lasso model.

The Lasso models with 50 iterations indicated that bridging the log(L/R) difference between TR-KA and the combined CT and TR-NK group would require a uniform increase of 19.61 units across the 5 fNIRS signals. Based on the descriptive statistics of sRMS in the combined CT and TR-NK group (standard deviations (SD) = 63.02, total physiological range of 613.11 with Min = −448.83, Max = 164.28), this equates to a shift of approximately 0.31 SD (equivalent to Cohen’s d of 0.31, which is a small to moderate effect size) or 3.20% of signals.

Since a 0.31 SD or 3.20% shift of signals falls well within the biologically observed range, these results suggest that it is mathematically plausible that differences in the strength of these 5 specific fNIRS signals in the PFC are sufficient to explain the higher log(L/R) in TR-KA than TR-NK and CT.

## DISCUSSION

In this study, we found that the baseline craving for a reinforcer at no cost (intensity of demand, Q0) was significantly larger in TR than in control subjects, regardless of the presence of kleptomania. However, TR-KA uniquely exhibited higher demand elasticity (α), reflecting profound intolerance to price changes during hypothetical purchases. Moreover, using the BART, TR-KA was found to have a significantly higher loss aversion than both TR-NK and CT. Notably, a correlation between the relative loss-to-risk aversion ratio measured in the BART and the elasticity in the HPT was observed in TR-KA, but not in TR-NK and CT. By inferring from PFC measurements in CT and TR-NK in which the relative loss-to-risk aversion ratio was found to be asymmetrically associated with the left and right PFC, such heightened loss aversion in TR-KA may be associated with an imbalance in left and right prefrontal cortical (PFC) activity.

Contrary to our initial hypothesis that theft recidivists might exhibit altered behavioral economic profiles resembling those of patients with addictive disorders, specifically increased risk-taking and reduced loss aversion (Canning et al., 2022; Genauck et al., 2017; Krmpotich et al., 2015; Thrailkill et al., 2022), this study found heightened loss aversion in theft recidivists with kleptomania. In addition, in the HPT, individuals with substance use disorders (SUD) typically display increased profiles across operant demand metrics, such as higher Q0 (desiring a large quantity of the substance), lower elasticity (ignoring price hikes), and a high maximum expenditure O_max_ (willingness to spend their last dollar to obtain it) (MacKillop, Goldenson, Kirkpatrick, & Leventhal, 2019; Zvorsky et al., 2019). While both TR, regardless of kleptomania, and SUD individuals share a high initial demand (Q0), suggesting that alterations may be partly overlapping between them, kleptomania is uniquely characterized by a high sensitivity to price hikes, which is the opposite of the low-price sensitivity seen in SUD. However, this does not exclude the possibility that kleptomania is not an addiction disorder, given that addiction disorder is characterized by concurrent impulsivity and compulsivity (Robbins, Gillan, Smith, de Wit, & Ersche, 2012), yet its expression of symptoms (e.g., SUD vs. behavioral addiction, which behavior to be addicted in behavioral addiction) may vary depending on what other mechanisms are involved (Goto, Krieger, Won, & Lee, 2026). Thus, whereas SUD is associated with lower monetary loss aversion and stronger reward-seeking desires (Krmpotich et al., 2015), individuals with kleptomania possess a stronger inherent aversion to monetary loss. In contrast, risk-taking propensity in TR, regardless of kleptomania, was not different from that of the control group.

The neural correlates of the relative loss-to-risk aversion ratio in this study suggest that the PFC plays an important role in economic decision-making. Our findings align with those of other studies in normal subjects, demonstrating that decision-making under uncertainty relies on an asymmetric division of labor between the hemispheres. Thus, the left and right PFC evaluate potential gains and losses, driving risk-averse and loss-averse behaviors, respectively (Huang et al., 2017; Ye et al., 2015). Accordingly, stronger left than right PFC activity is expected to increase risk-taking, whereas stronger right than left PFC activity is expected to drive greater loss aversion. Based on this framework, a fundamental imbalance between the left and right PFC may be associated with the pathophysiology of kleptomania.

This study had several major limitations. First, the study had a relatively small sample size and lacked racial and sociocultural diversities in participants, which limited the statistical power and generalizability of the results. Second, the study design was cross-sectional rather than longitudinal, precluding any definitive conclusions regarding the causality of the observed behavioral economic profiles. Moreover, due to logistical and environmental constraints, fNIRS measurements could not be directly conducted on participants diagnosed with kleptomania. Consequently, although this issue has been at least partly conquered by the in-silico perturbation analysis, which estimates approximately 3 to 4% increase and decrease of PFC activity may underlie heightened loss aversion in kleptomania, PFC activity profiles that may be associated with kleptomania had to be still inferred indirectly from the data of control subjects and theft recidivists without kleptomania.

The present study aimed to understand the underlying neurobehavioral mechanisms of theft recidivism with and without kleptomania could have some implications for society. Thus, redefining kleptomania not as a mere criminal action but as a quantifiable impairment in economic decision-making and loss evaluation would help reframe how society addresses this issue. Recognizing such neurobehavioral distinctions allows legal and healthcare systems to shift toward more specialized therapeutic interventions.

In conclusion, the findings of this study challenge the conventional view of theft as a reward-seeking behavior, revealing that recurrent stealing in kleptomania may be driven by a fear of monetary loss. Thus, recurrent stealing with kleptomania may stem from a fundamentally higher loss aversion that impairs cognitive computation during the trade-off between the products and their prices. This may be associated with an imbalance between left and right PFC activities, providing a novel perspective on the pathophysiology and therapeutic targets of intervention for kleptomania.

## CRediT author contributions statements

YG contributed to conceptualization, methodology, formal analysis, investigation, data curation, project administration, writing, visualization, validation, supervision, and funding acquisition. SA contributed to conceptualization, resources, and writing. CK contributed to conceptualization, resources, and writing. YAL contributed to formal analysis and writing. All authors have reviewed, edited, and approved the manuscript prior to submission.

## Acknowledgements

We would like to thank the staff of the Non-Profit Organization Kurashi O-en Network (Nagoya, Japan), Nishi Hongwanji Byakkoso (Kyoto, Japan), Non-Profit Organization Kyoto MAC (Kyoto, Japan), Kyoto Hogo Ikusei Kai (Kyoto, Japan), and Liberty Women’s House Olive (Otsu, Japan) for recruiting participants, scheduling surveys, and providing various technical supports.

