## Supplementary Materials for "Behavioral Economic Profiles in Theft Recidivists with and without Kleptomania"

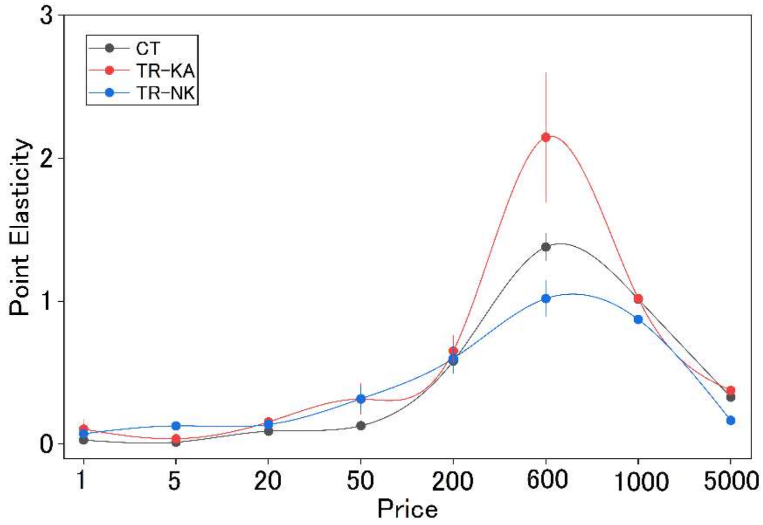


**Supplementary Figure S1.** Elasticity α was calculated by averaging the point elasticity at each price, defined as

$$E_p = \frac{dQ}{dP} \cdot \frac{P}{Q}$$

where dQ/dP is the derivative of quantity with respect to price, and P and Q are the specific price and quantity of snacks/sweets purchased at that exact price, respectively. The panel shows point plasticity at each price for control subjects with no criminal records (CT), theft recidivists with a diagnosis of kleptomania (TR-KA), and those without kleptomania (TR-NK).

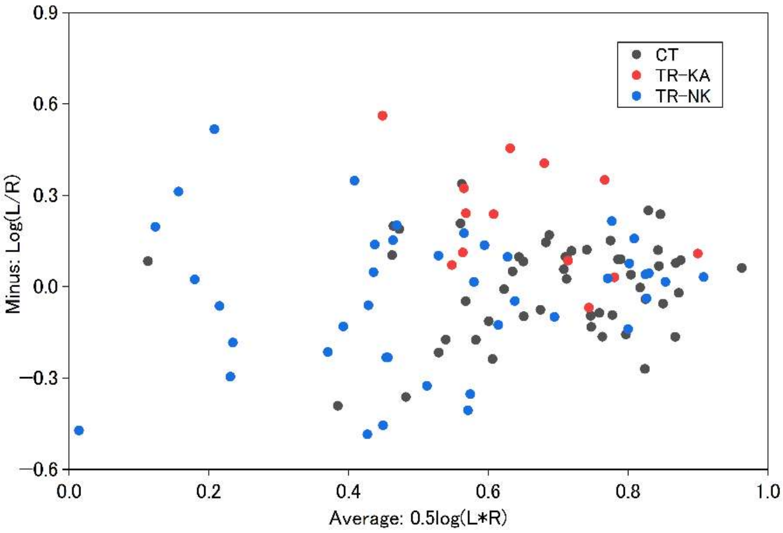


**Supplementary Figure S2.** The larger log(L/R) in TR-KA than in CT and TR-NK may be a consequence of a larger loss frame score (higher loss aversion), smaller gain frame score (higher risk aversion), or both; however, it was unclear which factor contributed more. This was evaluated using the minus vs. average (MA) plot, where the minus on the y-axis is the relative ratio log(L/R), and the average is the overall magnitude 0.5log(L•R). In this plot, a higher Minus in TR-KA than in CT and TR-NK, but a comparable Average of TR-KA relative to that of CT and TR-NK, were observed, suggesting that the larger log(L/R) in TR-KA is primarily due to the larger loss frame score (higher loss aversion).
